# Factors influencing maternal microchimerism throughout infancy and its impact on infant T cell immunity

**DOI:** 10.1101/2021.03.02.432825

**Authors:** Christina Balle, Blair Armistead, Agano Kiravu, Xiaochang Song, Anna-Ursula Happel, Angela Hoffmann, Sami B. Kanaan, J. Lee Nelson, Clive M. Gray, Heather B. Jaspan, Whitney E. Harrington

## Abstract

Determinants of the acquisition and maintenance of maternal microchimerism (MMc) during infancy and the impact of MMc on infant immune responses are unknown. We examined factors which influence MMc detection and level across infancy and the effect of MMc on T cell responses to BCG vaccination in a cohort of HIV exposed, uninfected and HIV unexposed infants in South Africa. MMc was measured in whole blood from 58 infants using a panel of quantitative PCR assays at day one and 7, 15, and 36 weeks of life. Infants received BCG at birth, and selected whole blood samples from infancy were stimulated in vitro with BCG and assessed for polyfunctional CD4+ T cell responses. MMc was present in most infants across infancy with levels ranging from 0-1,193/100,000 genomic equivalents and was positively impacted by absence of maternal HIV, maternal-infant HLA compatibility, infant female sex, and exclusive breastfeeding. Initiation of maternal antiretroviral therapy prior to pregnancy partially restored MMc level in HIV exposed, uninfected infants. Birth MMc was associated with an improved polyfunctional CD4+ T cell response to BCG. These data emphasize that both maternal and infant factors influence the level of MMc, which may subsequently impact infant T cell responses.

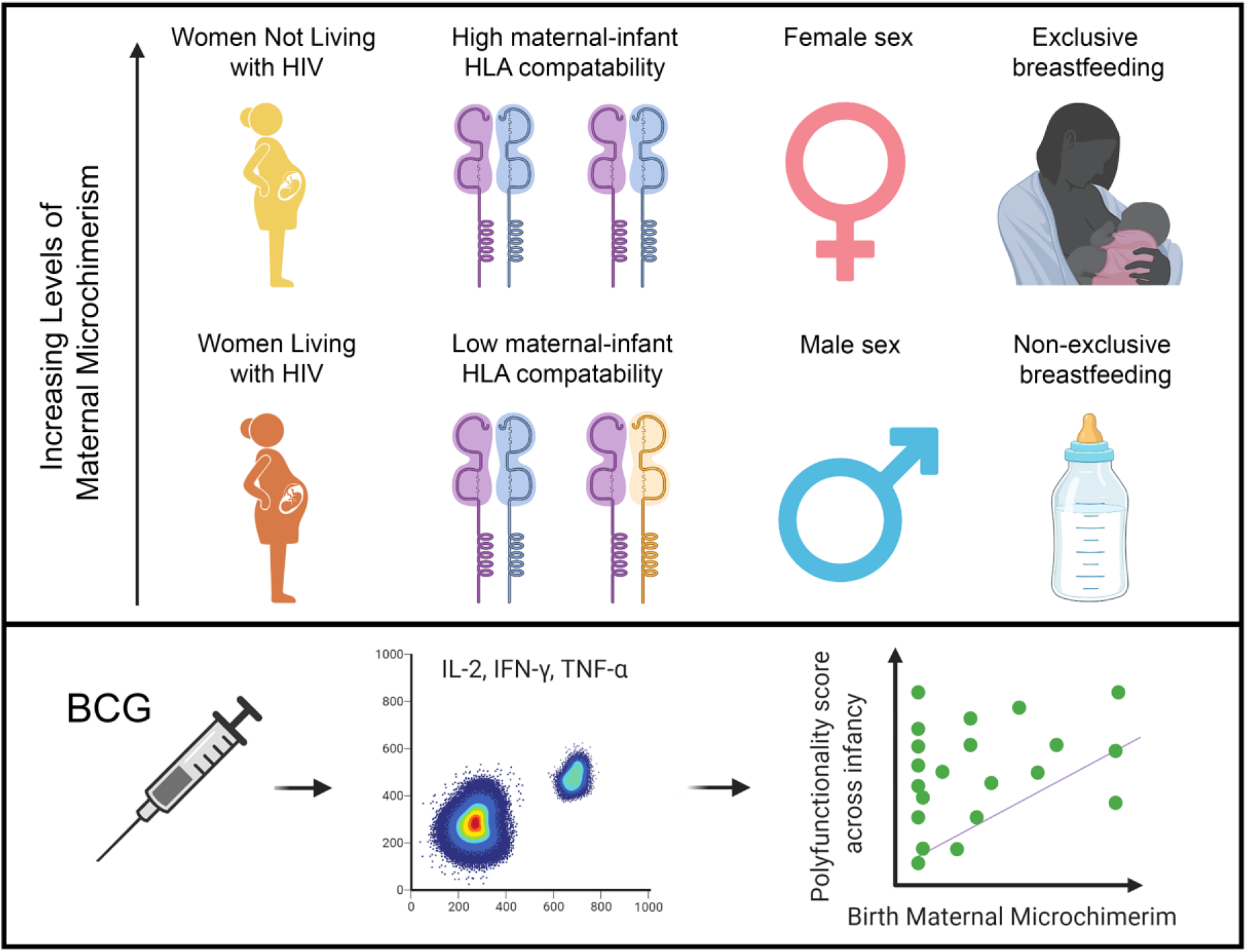

## Introduction

During pregnancy maternal cells are transferred to the fetus and persist in small quantities postnatally, a phenomenon known as maternal microchimerism (**MMc**). The impact of these maternal cells on the development of the fetal and infant immune system is largely unknown. MMc is acquired as early as the second trimester (1,2), is found in immune and nonimmune cells throughout the body (3–5), and is maintained into adulthood (3,6). However, there are limited data on factors that determine the quantity of cells a fetus acquires or those that impact the level of MMc across infancy. In a Tanzanian cohort, we have demonstrated that placental malaria infection, particularly inflammatory placental malaria, was associated with increased detection and level of cord blood MMc (7). In animal models, breastfeeding has been associated with increased maintenance of in utero acquired maternal cells, thought to be the product of increased tolerance to non-inherited maternal antigens (**NIMA**). Further, postnatal transfer of maternal cells to the offspring via breastfeeding has been demonstrated in rodents (12–15) and non-human primates (16), suggesting that the maternal graft in the infant may be a composite of cells acquired transplacentally and via the breastmilk.

Maternal HIV, even in the absence of vertical transmission, is associated with placental inflammation, including increased expression of inflammatory cell adhesion molecules, CD8 T cell infiltration (17), chorioamnionitis, and deciduitis (18). The impact of maternal HIV on MMc in the infant is unknown. Infants who are HIV exposed but uninfected (**iHEU**) experience increased morbidity and mortality compared to their HIV unexposed (**iHU**) counterparts (19). We have previously shown that iHEUs have altered T cell functionality to Bacillus Calmette-Guérin (**BCG**) and superantigens, including lower interferon (**IFN**)-γ and polyfunctional cytokine (IFN-γ, interleukin (**IL**)-2, and IL-17) responses (20), suggesting that altered immunity may contribute to their increased morbidity. Multiple potential mechanisms have been explored to explain such altered immunity, including differences in infant T cell repertoire (21) or changes to infant gut microbiome (22,23), secondary to altered in utero environment including antiretroviral therapy (**ART**) exposure, feeding modality, or maternal co-morbidities (24–27). While three studies have investigated the relationship between MMc, HIV, and the risk of vertical transmission (28–30), no studies to date have examined differences in MMc in iHEU versus iHU as a potential mediator of altered T cell responses in the offspring.

Maternal cells may shape fetal and infant immunity via at least three mechanisms. First, prior work has demonstrated an enrichment of maternal cells in antigen-experienced T cells from cord blood (31), suggesting that the offspring may acquire maternal antigen-specific T cells that could have direct effect when they encounter their cognate antigen in the infant. Second, the acquisition of MMc may indirectly shape the development of the fetal and infant immune system, including influencing the function of myeloid cells, as was recently demonstrated in mice (32). Third, MMc is associated with the development of NIMA-specific regulatory T cells (Tregs), which suppress effector T cell responses to maternal alloantigens (11). Encounter of neoantigen in the presence of such Tregs may induce cross-tolerance as has been described in transplant biology (8,33,34). Supporting the potential for MMc to impact infant outcomes, we have previously found that detection of cord blood MMc in Tanzanian infants was associated with decreased risk of symptomatic malaria (7), and whole blood MMc detection in American children was found to be protective from the later development of asthma (35).

Here, we investigate factors associated with the acquisition and maintenance of MMc in South African infants including the potential impact of maternal HIV, maternal-infant HLA compatibility, infant sex, and mode of feeding. We further assess the impact of MMc on infant T cell responses directed against BCG, a common neonatal antigen exposure.

## Results

### Cohort characteristics

We screened 90 mother-infant pairs for a maternal-specific marker. Of these, 58 mothers had a unique marker which could be targeted to detect MMc in the background of the infant. There were no differences in the characteristics of the mother-infant pairs that could or could not be targeted for measurement of MMc (**Supplementary Table 1**). The mean maternal age at delivery was 27 years (standard deviation (**sd**) 5.1) (**Table 1**). This was the first pregnancy for 20.7% of the mothers (12/58) and the first delivery for 34.5% (20/58). The mean estimated gestational age (**EGA**) at delivery was 39.4 weeks (sd 1.37), and mean infant birth weight was 3,207 grams (sd 416). Just under half of the infants were female (26/58, 44.8%). Nearly two-thirds of the mothers were living with HIV (37/58, 63.8%), of whom just over half (20/37, 54.1%) had received ART prior to conception (**early ART**) while the remaining mothers initiated ART during pregnancy (**late ART**) at a mean gestational age of 21 weeks (sd 8.3).

**Table 1.** Cohort characteristics.

|  | **Total**<br>**(n=58)** |
| --- | --- |
| **Maternal age years, mean (sd)** | 27 (5.1) |
| **Gravidity, n (%)** |  |
| Primigravidae | 12 (20.7) |
| Secondigravidae | 25 (43.1) |
| Multigravidae | 21 (36.2) |
| **Infant sex, female, n (%)** | 26 (44.8) |
| **Birth weight, grams, mean (sd)** | 3,207 (416) |
| **Gestational age at delivery, weeks, mean (sd)** | 39.4 (1.37) |
| **Delivery method, vaginal, n (%)** | 58 (100) |
| **HIV exposed, n (%)** | 37 (63.8) |
| **Maternal antiretroviral therapy** |  |
| Prior to pregnancy, n (%) | 20/37 (54.1) |
| Initiated during pregnancy, n (%) | 17/37 (45.9) |
| Gestational age at initiation, weeks, mean (sd) | 21 (8.3) |
| **BCG strain used for vaccination** |  |
| BCG-Denmark, n (%) | 39 (67.2) |
| BCG-Russia, n (%) | 19 (32.8) |

### Maternal microchimerism at birth is modified by HIV exposure, HLA compatibility, and infant sex

The mean genomic equivalents (**gEq**) analysed in whole blood samples collected at day one of life (henceforth referred to as birth) was 148,187 (sd 52,867) per sample. Two outcomes were considered for each analysis: 1) the detection of any MMc (no or yes) and 2) the level of MMc. At birth, MMc was detectable in 23/58 (39.7%) infants with levels ranging from 0-24/100,000 gEq (**Figure 1A-B**). In order to identity factors associated with the detection and quantity of maternal cells, we evaluated the potential impact of maternal HIV status, gravidity, maternal-infant HLA Class II compatibility (henceforth referred to as HLA compatibility), EGA, and infant sex. In univariate models, none of the covariates were significantly associated with increased detection of MMc (**Figure 2, Table 2**). However, infant female sex was associated with significantly higher level of MMc (unadjusted detection rate ratio (**DRR**) 3.71 [95% CI 1.11-12.4], P=0.033) (**Figure 2F, Table 2**), and there were non-significantly higher MMc levels in infants with increasing HLA compatibility and lower MMc levels in iHEU and in infants of multigravidae (**Figure 2, Table 2**). In multivariate models including maternal HIV status, gravidity, HLA compatibility, and infant sex, the association between HIV exposure and level of MMc at birth became significant (adjusted DRR 0.34 [95% CI 0.13-0.89], P=0.028), as did the association with HLA compatibility (adjusted DRR 1.96 [1.16-3.32], P=0.017), while the impact of infant sex remained significant (adjusted DRR 3.36 [95% CI 1.08-10.5], P=0.037) (**Table 2**).

**Figure 1:**
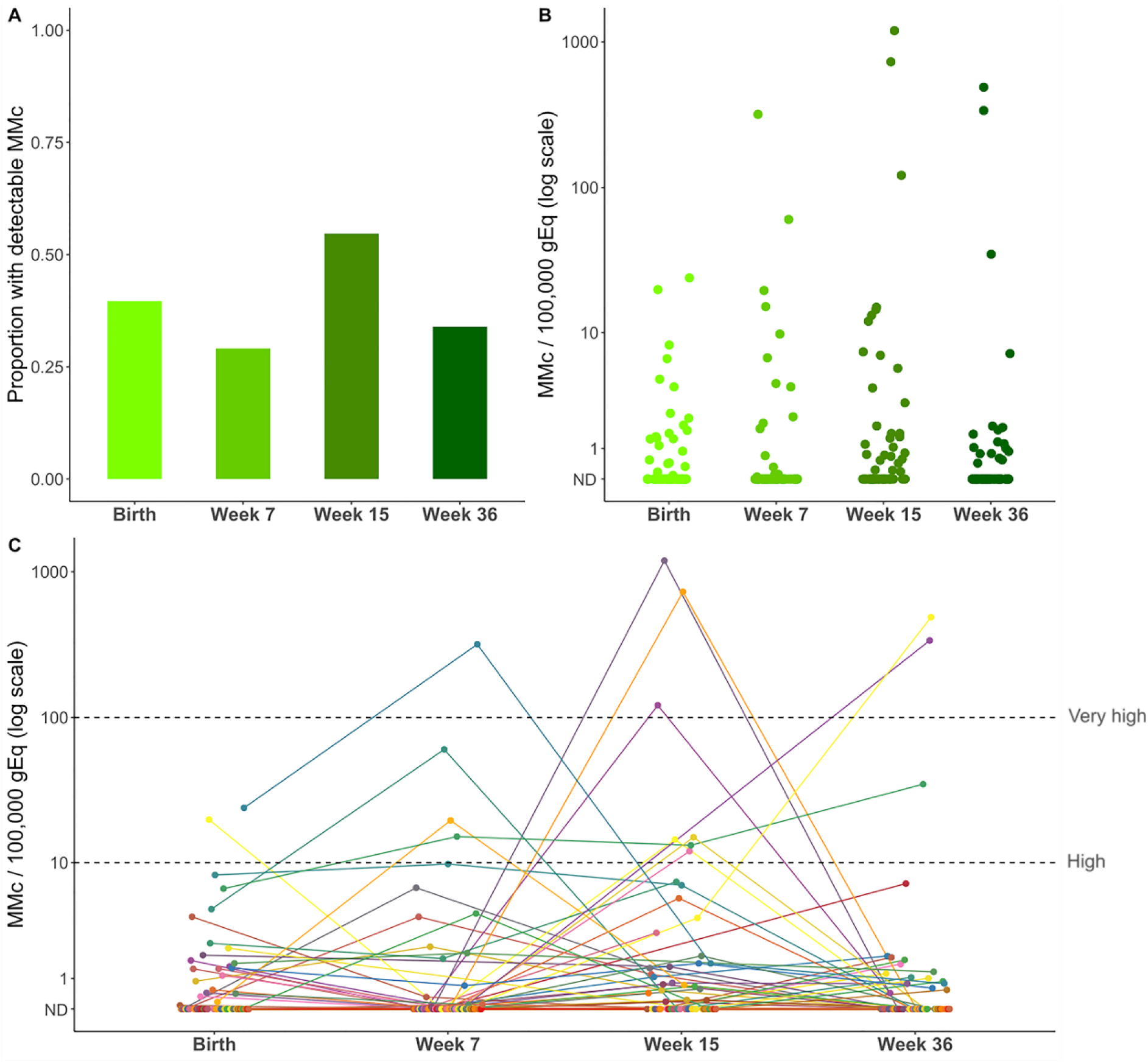
Maternal microchimerism is dynamic and persist throughout infancy. Maternal microchimerism (MMc) expressed as microchimeric equivalents per infant genomic equivalents (gEq) assessed. Not detected (ND). (A) Proportion of infants with detectable MMc at birth (N=58), week 7 (N=55), week 15 (N=53), and week 36 (N=53) of life. (B) Quantitative levels of MMc across infancy. (C) Within infant MMc kinetics. The dotted lines highlight MMc levels above 10/100,000 gEq (High) and above 100/100,000 gEq (Very high).

**Figure 2.**
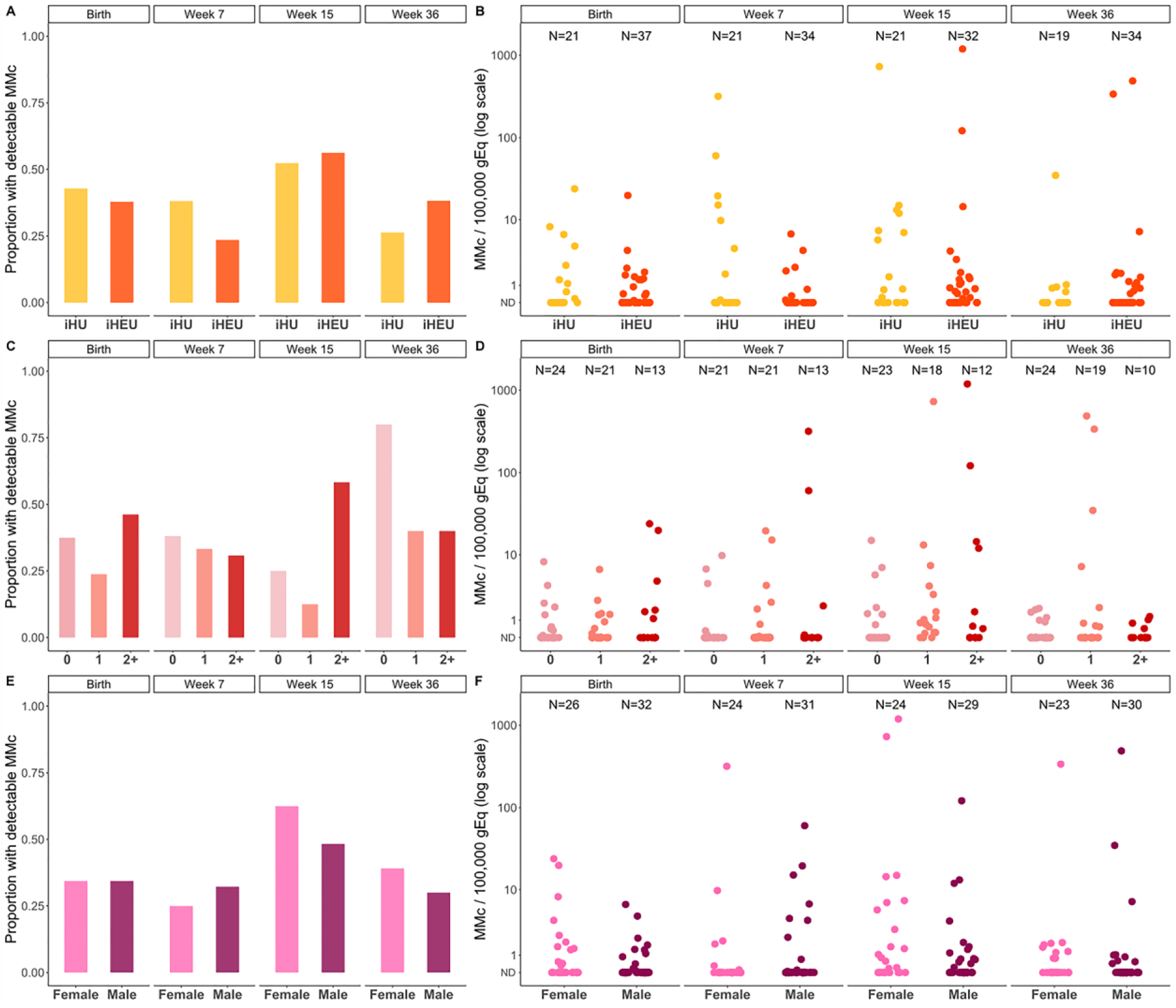
Maternal microchimerism across infancy is associated with HIV exposure, maternal-infant HLA Class II compatibility, and infant sex. Maternal microchimerism (MMc) expressed as microchimeric equivalents per infant genomic equivalents (gEq) assessed. Not detected (ND), HIV unexposed infants (iHU), HIV exposed, uninfected infants (iHEU). Detection of MMc in infants at day one, week 7, week 15 and week 36 of life by (A) HIV exposure, (C) maternal-infant HLA compatability, and (E) infant sex. Level of MMc across infancy by (B) HIV exposure, (D) maternal-infant HLA compatability, and (F) infant sex.

**Table 2.** Association between covariates and detection and level of maternal microchimerism at birth.

|  | Detection of MMc at birth |  |  |  |
| --- | --- | --- | --- | --- |
|  | OR (95% CI) | P value | Adj. OR (95% CI) | Adj. P value |
| **Maternal HIV** | 0.82 (0.27-2.49) | 0.720 | 0.79 (0.25-2.46) | 0.690 |
| **HLA Compatibility** | 1.30 (0.76-2.22) | 0.344 | 1.30 (0.73-2.32) | 0.369 |
| **Gravidity** |  |  |  |  |
| Primigravidae | REF |  | REF | - |
| Secondigravidae | 0.95 (0.22-4.12) | 0.943 | 1.03 (0.24-4.47) | 0.970 |
| Multigravidae | 0.88 (0.19-4.05) | 0.867 | 1.15 (0.25-5.28) | 0.859 |
| **EGA** | 0.96 (0.58-1.59) | 0.869 | - | - |
| **Infant sex, female** | 1.65 (0.55-4.94) | 0.368 | 1.65 (0.54-5.01) | 0.377 |
|  | Level of MMc at birth |  |  |  |
|  | DRR (95% CI) | P value | Adj. DRR (95% CI) | Adj. P value |
| **Maternal HIV** | 0.45 (0.11-1.92) | 0.282 | 0.34 (0.13-0.89) | **0.028** |
| **HLA Compatibility** | 1.94 (0.94-4.01) | 0.073 | 1.96 (1.16-3.32) | **0.017** |
| **Gravidity** |  |  |  |  |
| Primigravidae | REF |  | REF | - |
| Secondigravidae | 0.65 (0.12-3.59) | 0.622 | 0.49 (0.11-2.17) | 0.350 |
| Multigravidae | 0.27 (0.06-1.17) | 0.081 | 1.01 (0.28-3.65) | 0.990 |
| **EGA** | 0.96 (0.58-1.58) | 0.867 | - | - |
| **Infant sex, female** | 3.71 (1.11-12.4) | **0.033** | 3.36 (1.08-10.5) | **0.037** |
| Detection of maternal microchimerism (MMc) at birth was analyzed using logistic regression controlling for total genomic equivalents (gEq) assessed. Level of MMc at birth was analyzed using negative binomial regression accounting for both microchimeric and total gEq. Adjusted models included maternal-infant HLA Class II score, gravidity, and infant sex. Adj: adjusted, EGA: estimated gestational age, OR: odds ratio, DRR: detection rate ratio. |  |  |  |  |

### Maternal microchimerism across infancy is dynamic and modified by HIV exposure, HLA compatibility, infant sex, and mode of feeding

In addition to the birth time point, we also evaluated MMc at week 7, 15 and 36 of life. Across infancy, MMc was detectable at any time point in 44/58 (75.9%) infants with a range of detection of 1-3 timepoints and levels ranging from 0-1,193/100,000 gEq (**Figure 1A-B**). Within individuals, the level of MMc detected across infancy was dynamic (**Figure 1C**). Levels of MMc higher than typically detected (>10/100,000 gEq) were identified at at least one time point in 12/58 (20.7%) infants, whereas very high MMc (>100/100,000 gEq) was identified at week 7 in one infant (318/100,000 gEq), at week 15 in three infants (121, 729, and 1,193 per 100,000 gEq), and at week 36 in two infants (338 and 488 per 100,000 gEq) (**Figure 1C**). Across the group, MMc level increased between birth and 15 weeks, then fell at 36 weeks (**Figure 1A-B**).

We assessed whether the covariates associated with MMc at birth, as well as the additional covariates of feeding modality and age of the infant were associated with detection and level of MMc across infancy. Only the degree of HLA compatability was significantly associated with increased detection of MMc (adjusted OR 1.54 [95% CI 1.10-2.15], P=0.012) (**Figure 2**). However, HIV exposure was associated with a significantly lower level of MMc (adjusted DRR: 0.37 [95% CI: 0.15-0.94], P=0.036) (**Figure 2B**, **Table 3**). Higher HLA compatibility was associated with increased level of MMc (adjusted DRR 3.63 [95% CI 2.45-5.39], P<0.001) (**Figure 2D**) (**Table 3**). Compared to male infants, female infants had significantly higher levels of MMc throughout infancy (adjusted DRR 4.56 [95% CI 1.38-15.1], P=0.013) (**Figure 2E-F, Table 3**). Exclusive breastfeeding (**EBF**) was associated with non-significantly higher levels of MMc compared to non-EBF (**NBF**) (adjusted DRR 4.05 [95% CI 0.85-19.4], P=0.080) (**Table 3**). By week 36, only one infant remained exclusively breastfed, so to further confirm that the associations by mode of feeding were not driven by this skewed distribution, we re-ran our multivariate models excluding week 36 from the analysis. Exclusive breastfeeding was associated with significantly higher levels of infant MMc in the unadjusted analysis (unadjusted DRR 11.5 [95% CI 3.13-42.3], P<0.001) and non-significantly higher levels in the adjusted model (adjusted DRR 2.73 [95% 0.92-8.84], P=0.068, **Supplementary Table 2; Figure 3**). Finally, older age was significantly associated with higher level of MMc (adjusted DRR per week: 1.13 [95% CI 1.07-1.20], P=<0.001) (**Figure 1B**) (**Table 3**).

**Figure 3.**
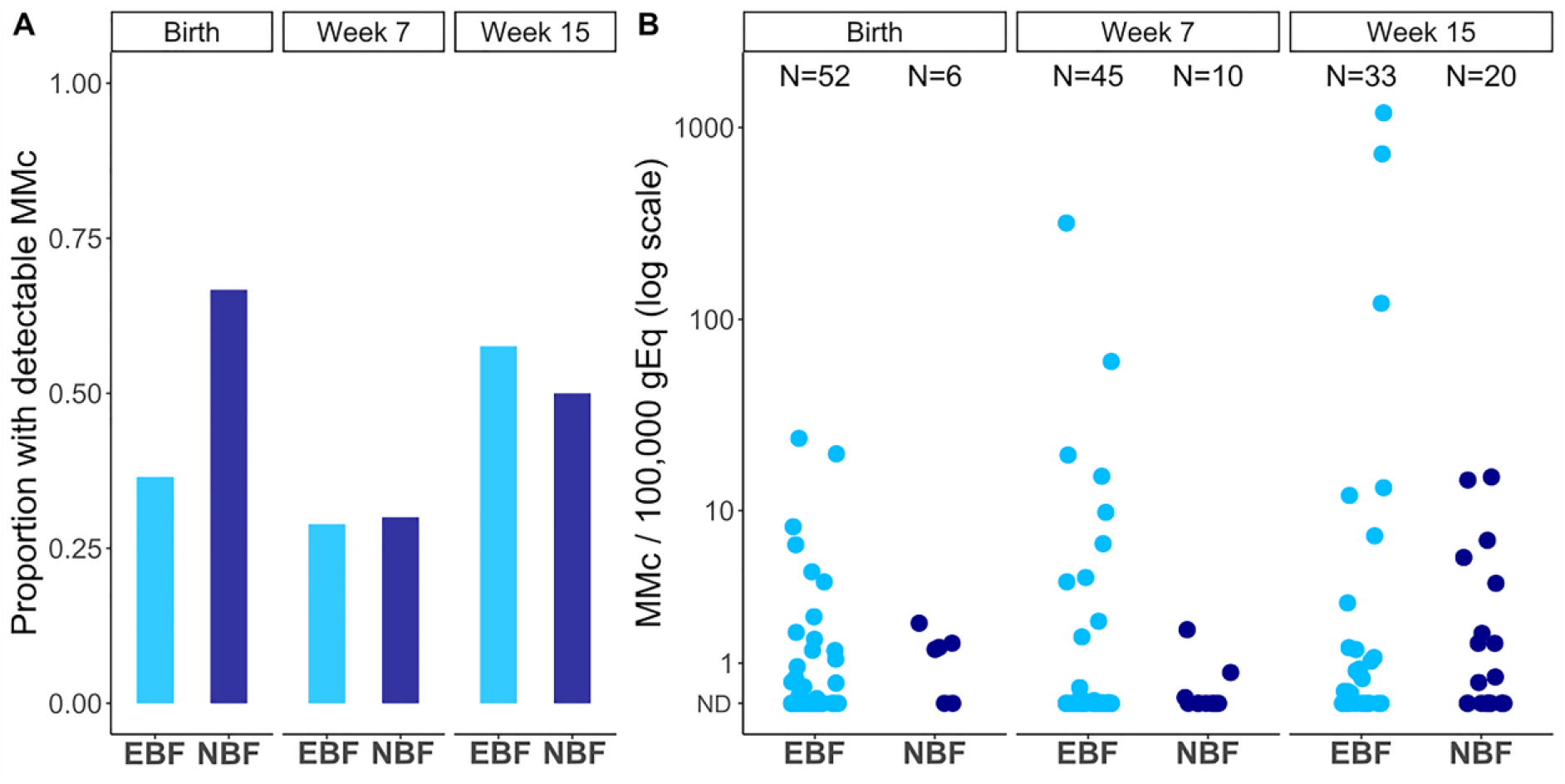
Maternal microchimerism across infancy is associated with mode of feeding. Maternal microchimerism (MMc) expressed as microchimeric equivalents per infant genomic equivalents (gEq) assessed. Not detected (ND), exclusively breastfed (EBF), non-exclusively breastfed (NBF). (A) Detection of MMc in infants at day one, week 7 and week 15 of life by mode of feeding. (B) Level of MMc across infancy by mode of feeding.

**Table 3.** Association between covariates and detection and level of maternal microchimerism across infancy.

|  | **Detection of MMc across infancy** |  |  |  |
| --- | --- | --- | --- | --- |
|  | **OR (95% CI)** | **P value** | **Adj. OR (95% CI)** | **Adj. P value** |
| **Maternal HIV** | 0.91 (0.44-1.89) | 0.802 | 0.83 (0.40-1.72) | 0.612 |
| **HLA Compatibility** | 1.41 (1.00-1.99) | **0.048** | 1.54 (1.10-2.15) | **0.012** |
| **Gravidity** |  |  |  |  |
| Primigravidae | REF | - | REF | - |
| Secondigravidae | 1.47 (0.55-3.90) | 0.439 | 1.38 (0.45-4.20) | 0.571 |
| Multigravidae | 1.80 (0.65-4.98) | 0.260 | 2.17 (0.72-6.58) | 0.169 |
| **Infant sex, female** | 1.35 (0.69-2.66) | 0.385 | 1.51 (0.71-3.20) | 0.286 |
| **Exclusively breastfed** | 0.91 (0.49-1.68) | 0.754 | 0.94 (0.40-2.21) | 0.889 |
| **Infant age, weeks** | 1.00 (0.98-1.02) | 0.964 | 1.00 (0.98-1.03) | 0.789 |
|  | **Level of MMc across infancy** |  |  |  |
|  | **DRR (95% CI)** | **P value** | **Adj. DRR (95% CI)** | **Adj. P value** |
| **Maternal HIV** | 0.91 (0.17-4.91) | 0.913 | 0.37 (0.15-0.94) | **0.036** |
| **HLA Compatibility** | 5.05 (3.60-7.07) | **<0.001** | 3.63 (2.45-5.39) | **<0.001** |
| **Gravidity** |  |  |  |  |
| Primigravidae | REF | - | REF | - |
| Secondigravidae | 11.2 (2.75-45.7) | **0.001** | 2.04 (0.44-9.52) | 0.365 |
| Multigravidae | 0.74 (0.14-4.04) | 0.730 | 1.91 (0.44-8.19) | 0.449 |
| **Infant sex, female** | 8.93 (2.43-32.8) | **0.001** | 4.56 (1.38-15.1) | **0.013** |
| **Exclusively breastfed** | 2.52 (0.54-11.7) | 0.238 | 4.05 (0.85-19.4) | 0.080 |
| **Infant age, weeks** | 1.05 (1.03-1.07) | **<0.001** | 1.13 (1.07-1.20) | **<0.001** |
| Detection of maternal microchimerism (MMc) across infancy was analyzed using generalized estimating equation (GEE) models with binomial distribution, controlling for |  |  |  |  |

total genomic equivalents (gEq) assessed. Level of MMc across infancy was analyzed using GEE models with negative binomial distribution accounting for both microchimeric and total gEq. Adjusted models included maternal-infant HLA Class II score, mode of feeding, gravidity, and infant sex. Adj.: adjusted, EGA: estimated gestational age, DRR: detection rate ratio, OR: odds ratio.

### Initiation of ART prior to conception partially restores maternal microchimerism in iHEU

We next assessed whether the timing of ART initiation (pre-conception or during gestation) altered the effect of maternal HIV on MMc by considering the iHEUs by timing of maternal ART initiation (early or late ART) and repeated the analyses. While both of the HIV-exposed groups had lower levels of MMc compared to the iHUs at birth, this was only significant for infants born to mothers with late ART initiation (**Table 4**). Similarly, late ART iHEUs had significantly lower level of MMc (adjusted DRR 0.14 [95% CI 0.03-0.59], P=0.007) (**Figure 4B, Table 4**) across infancy as compared to iHUs, while early ART iHEUs had similar MMc levels to the iHU infants (**Figure 4, Table 4**). When the two groups of iHEU were compared to each other, there was no difference in detection or level of birth MMc (adjusted OR=1.09, P=0.899, adjusted DRR= 0.20, P=0.772). However, there was an increased level (adjusted DRR=1.57, P=0.050) of MMc amongst early versus late ART iHEUs across infancy. Collectively, these data indicate that longer duration of maternal ART (from pre-conception) may partially restore MMc levels in iHEU.

**Figure 4.**
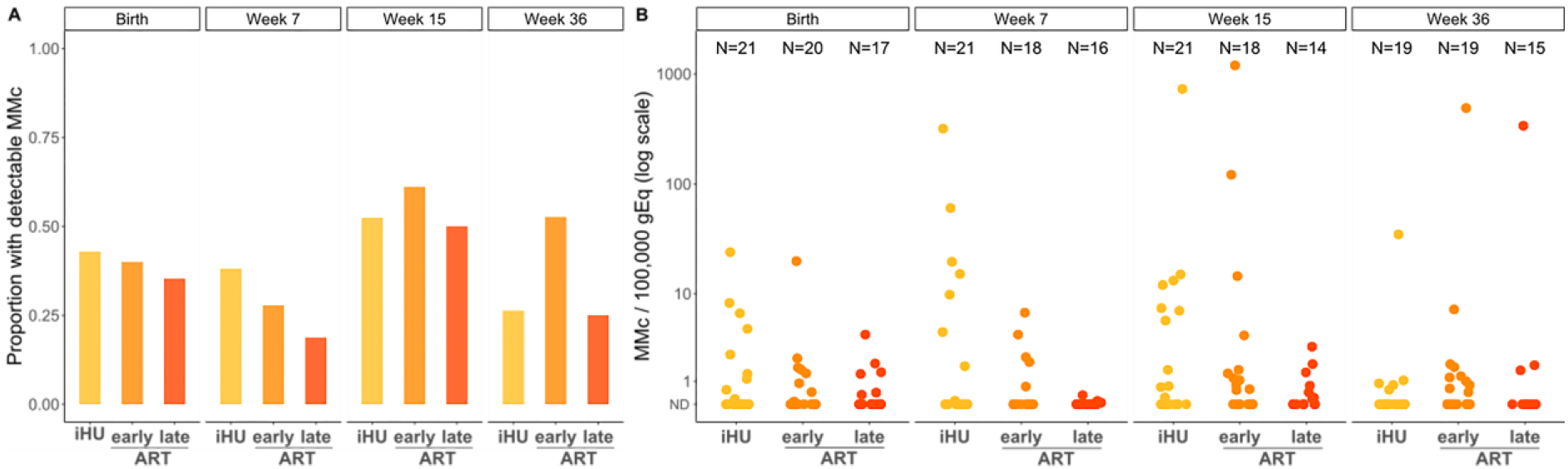
Maternal microchimerism across infancy is associated with duration of maternal antiretroviral therapy (ART). Maternal microchimerism (MMc) expressed as microchimeric equivalents per infant genomic equivalents (gEq) assessed. Not detected (ND), HIV unexposed infants (iHU), HIV exposed, uninfected infants whose mothers were on antiretroviral therapy (ART) pre-conception (early ART), HIV exposed, uninfected infants whose mothers initiated ART during pregnancy (late ART). (A) Detection of MMc in infants at day one, week 7, week 15 and week 36 of life by HIV exposure and duration of ART. (B) Level of MMc across infancy by HIV exposure and duration of ART.

**Table 4.** Association between HIV exposure and duration of antiretroviral therapy on detection and level of maternal microchimerism at birth and across infancy.

|  | **Detectable MMc at birth** |  |  |  |
| --- | --- | --- | --- | --- |
|  | **OR (95% CI)** | **P value** | **Adj. OR (95% CI)** | **Adj. P value** |
| **HIV unexposed** | REF | - | REF | - |
| **Early ART** | 0.89 (0.25- 3.10) | 0.852 | 0.83 (0.22-3.07) | 0.777 |
| **Late ART** | 0.73 (0.19-2.74) | 0.646 | 0.76 (0.19-2.93) | 0.685 |
|  | **Level of MMc at birth** |  |  |  |
|  | **DRR (95% CI)** | **P value** | **Adj. DRR (95% CI)** | **Adj. P value** |
| **HIV unexposed** | REF | - | REF | - |
| **Early ART** | 0.63 (0.12-3.25) | 0.577 | 0.37 (0.12-1.12) | 0.080 |
| **Late ART** | 0.29 (0.06-0.90) | **0.034** | 0.30 (0.09-1.02) | 0.053 |
|  | **Detectable MMc across infancy** |  |  |  |
|  | **OR (95% CI)** | **P value** | **Adj. OR (95% CI)** | **Adj. P value** |
| **HIV unexposed** | REF | - | REF | - |
| **Early ART** | 1.21 (0.54-2.71) | 0.649 | 1.03 (0.47-2.25) | 0.943 |
| **Late ART** | 0.63 (0.27-1.48) | 0.290 | 0.62 (0.25-1.50) | 0.287 |
|  | **Level of MMc across infancy** |  |  |  |
|  | **DRR (95% CI)** | **P value** | **Adj. DRR (95% CI)** | **Adj. P value** |
| **HIV unexposed** | REF | - | REF | - |
| **Early ART** | 1.44 (0.24-8.59) | 0.691 | 0.68 (0.25-1.86) | 0.454 |
| **Late ART** | 0.28 (0.03-2.43) | 0.250 | 0.14 (0.03-0.59) | **0.007** |
| Detection of maternal microchimerism (MMc) at birth was analyzed using logistic regression controlling for total genomic equivalents (gEq) assessed. Level of MMc at birth was analyzed using negative binomial regression. Adjusted birth models included maternal-infant HLA Class II score, gravidity, and infant sex. Detection of MMc across infancy was analyzed using generalized estimating equation (GEE) models with binomial distribution, controlling for total gEq assessed. Level of MMc across infancy was analyzed using GEE models with negative binomial distribution. Adjusted infant models included gravidity, infant sex, maternal-infant HLA Class II scoring, mode of feeding, and infant age. Adj: adjusted, ART: antiretroviral therapy, EGA: estimated gestational age, OR: odds ratio, DRR: detection rate ratio. |  |  |  |  |

### Maternal microchimerism at birth is associated with polyfunctional CD4+ T cell responses to BCG

Using previously measured BCG vaccine response data from this cohort (36), we sought to determine whether the detection or level of MMc at time of BCG vaccination (birth) would predict BCG-specific T cell responses during early infancy (week 7 and week 15). We used the COMPASS polyfunctional score (**PFS**), a clinically validated tool for assessing polyfunctional T cell function (37) to measure CD4+ T cell responses in 33 infants across 48 samples and adjusted for BCG strain (Danish versus Russian), HIV exposure, and infant age in our analyses. Both detection (adjusted coefficient: 0.340, P<0.001) (**Figure 5A**) and level of MMc (adjusted coefficient per 10/100,000 gEq: 0.272, P<0.001) (**Figure 5B**) at birth were positively associated with the PFS. In contrast, concurrent detection or level of MMc in the infant blood at week 7 or 15 was not associated with polyfunctionality of BCG response at the time point. (**Figure 5C,D**). Similar associations were found for the functionality score (**FS**) (**Supplementary Figure 1**).

**Figure 5.**
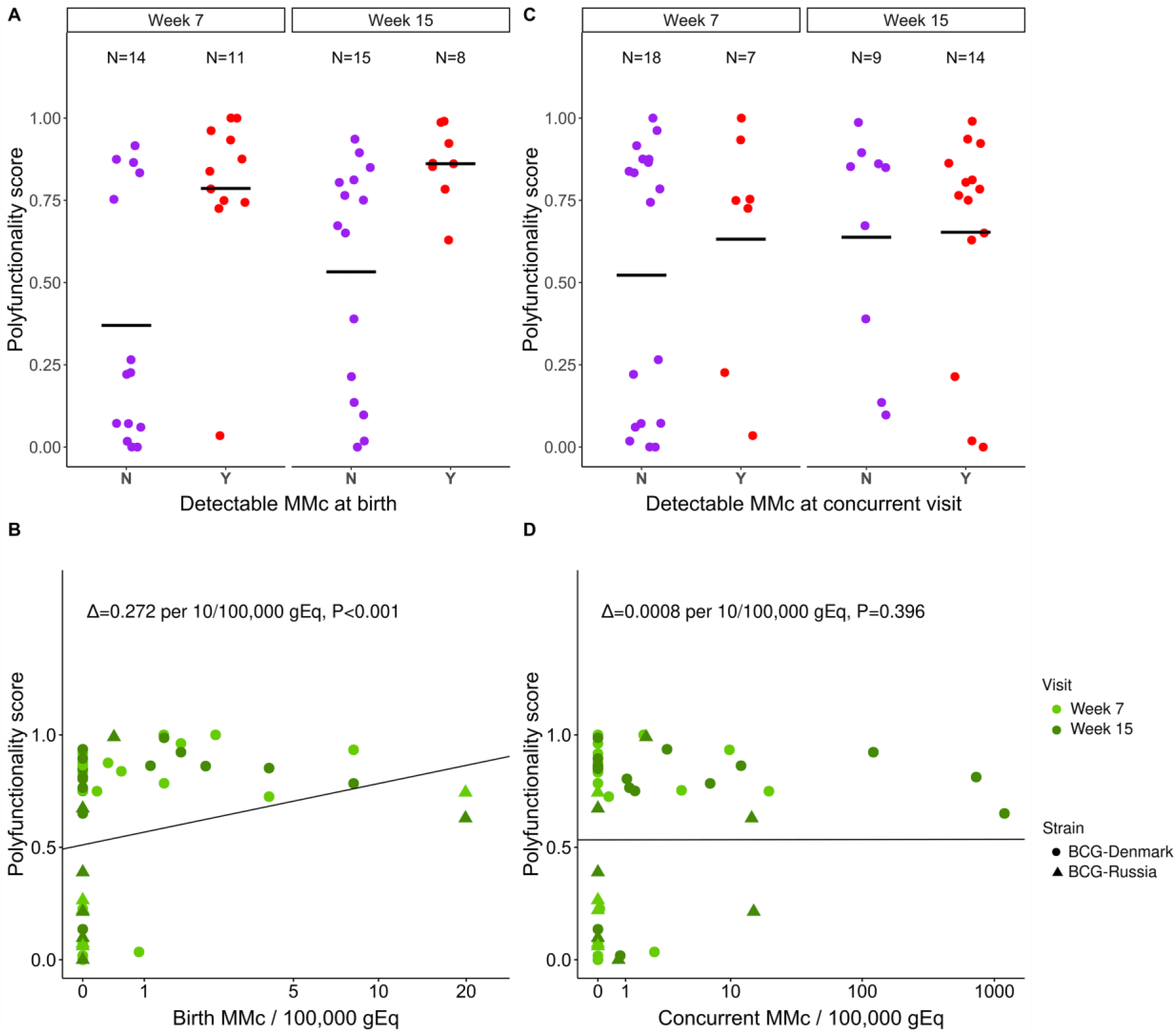
Maternal microchimerism at birth is positively associated with T cell responses to BCG vaccination. (A) Polyfunctionality score (PFS) at week 7 and week 15 of life by detection of maternal microchimerism (MMc) at birth (N=No (purple), Y=Yes (red)). (B) Association between PFS at week 7 (light green) and week 15 (green) of life and level of MMc per 100,000 genomic equivalents (gEq) at birth. Delta represents the adjusted effect size per 10/100,000 gEq. (C) PFS at week 7 and week 15 of life by detection of MMc at the concurrent time point (Y=Yes (red), N=No (purple)). (D) Association between PFS at week 7 (light green) and week 15 (green) of life and level of MMc per 100,000 genomic equivalents (gEq) at the concurrent time point. Delta represents the adjusted effect size per 10/100,000 gEq. Horizontal black lines indicate mean values.

To evaluate whether maternal BCG-responsive T cells directly contributed to the polyfunctional T cell responses to BCG detected in infants, we identified infants who had detectable MMc at birth, a PFS>0.6, and available peripheral blood mononuclear cells (**PBMC**) for functional evaluation (n=4). Infant PBMCs from these timepoints were stimulated with BCG culture filtrate proteins, followed by intracellular cytokine staining. T cells were sorted into polyfunctional, monofunctional, or non-responding populations, and we measured MMc level in each population by qPCR. While we were able to detect maternal T cells in sorted populations from one iHU, they remained the minority of the responding population, and no MMc was detected in the polyfunctional cells in the other three infants (**Table 5**), indicating that maternal BCG-specific T cells did not directly contribute to the polyfunctional T cell response to BCG in these infants.

**Table 5.** Level of MMc in sorted T cell populations following stimulation of infant PBMC with BCG culture filtrate proteins.

| Infant | MMc level |  |  |
| --- | --- | --- | --- |
|  | Polyfunctional T cells | Monofunctional T cells | Non-responsive T cells |
| 1 | 384/100,000 | 123/100,000 | 11/100,000 |
| 2 | 0 | 0 | 0 |
| 3 | 0 | 0 | 0 |
| 4 | 0 | 0 | 0 |

Available PBMC from infants with detectable birth MMc and COMPASS PFS>0.6 were stimulated with BCG culture filtrate proteins, analyzed for expression of IL-2, TNF-α , and IFN-γ by intracellular cytokine staining, and sorted into polyfunctional (two or more cytokines), monofunctional (one cytokine), and non-responsive (no cytokines) T cell groups. The MMc level in each sorted T cell population was determined by qPCR targeting a maternal specific allele.

## Discussion

This study is the first to measure and compare MMc acquisition and maintenance in iHEU and iHU during the first year of life. We observed MMc at least at one time point in the majority of infants in the cohort, with high and very high levels observed in some infants at the later time points. At birth, lack of HIV exposure, increased HLA compatibility, and female infant sex were positively associated with MMc. Across infancy, MMc increased up to a peak at 15 weeks of age and was positively associated with lack of HIV exposure, increased HLA compatibility, female infant sex, and exclusive breastfeeding. Finally, MMc at birth was associated with increased polyfunctionality to BCG later in life, suggesting that these maternal cells may have functional consequence for infant immune responses.

We observed a lower level of MMc in iHEU compared to iHU in this cohort. Human studies have found the highest proportion of MMc to be T cells both in fetal lymph nodes (11) and in cord blood from term pregnancies (31). Similarly in mice, the majority of breastmilk cells undergoing trans-epithelial migration were CD4+ and CD8+ T cells (14). Thus, we hypothesize that the reduction in MMc observed in iHEUs may be due to a reduction of CD4+ T cells in the mothers, leading to an overall reduced trafficking of these cells to the fetus and infant. Alternatively, HIV is associated with systemic immune dysregulation, chronic antigen presenting cell (**APC**) activation (38,39), and placental inflammation (17,18), which may lead to altered maternal-fetal tolerance and increased rejection of the maternal graft. The reduced MMc conferred by HIV exposure may contribute to abnormal immune responses and increased infectious morbidity in exposed offspring (19,40,41).

Interestingly, we observed that iHEUs born to mothers who initiated ART pre-conception had higher levels of MMc than those born to mothers who initiated ART during the current pregnancy, and that in the former, levels were comparable to those in iHUs. Early ART may normalize levels of MMc to those seen in iHU through the restoration of maternal CD4+ T cells and/or improvement in systemic immune dysregulation or placental inflammation. These potential mechanisms should be examined in future studies. As MMc acquisition has been observed as early as the beginning of the second trimester (1,2), initiating ARTs during a later antenatal visit may fail to restore normal MMc levels in the offspring.

In this cohort, HLA Class II compatibility between mother and infant was associated with increased levels of MMc. This observation is consistent with earlier work finding an association between MMc and maternal compatibility at the HLA class II DRB1 and DQB1 loci (42) as well as animal models (43). HLA play a critical role immune regulation (44,45) and increasing degree of maternal-infant HLA compatibility may lead to increased tolerance of the maternal graft.

There are limited data comparing MMc in female and male offspring. Here, we found that female infants had higher levels of MMc than male infants. This observation is consistent with prior studies comparing children aged one to seven years born to asthmatic and non-asthmatic mothers where MMc detection was higher in daughters (24.3%) compared to sons (16.9%) (28). Female offspring are protected from pregnancy loss under conditions of maternal stress relative to male offspring, suggesting a higher degree of maternal-fetal tolerance (46) where such tolerance may allow the microchimeric graft to persist in the offspring. Increased accumulation of MMc in female compared with male offspring may offer protection against complications in next-generation pregnancies as shown in mice (47). Further, differences in MMc levels between females and males during early infancy could potentially lead to distinct effects on infant immune development, including differences in vaccine responses and disease susceptibility (40). Additional studies are needed to elucidate the potential mechanism behind this finding.

Our results suggest that exclusive breastfeeding may be associated with higher levels of MMc across infancy compared to non-exclusive breastfeeding. Breastfeeding plays a major role in protection from infection in infants through transfer of antibody and other immunomodulatory components (48–51). Recent work has demonstrated that breastmilk derived IgA can mediate cross-generational impact on Treg development in the offspring, emphasizing the potential for non-genetically encoded heritability (52). Recent data from animal models has demonstrated that breastmilk-derived maternal cells (13–16), including pathogen-experienced T cells (12), can be detected in the offspring. Our study raises the possibility that postnatal MMc may be acquired via breastmilk in humans (53), which may be more pronounced during the early stages of lactation when breastmilk cells are more abundant (54) and infant gut permeability the highest (55). Alternatively, the increased levels of MMc in exclusively breastfed infants could be due to increased NIMA-specific tolerance leading to maintenance of in utero acquired MMc (8,9,11,47).

We observed that at the population level MMc increased with infant age, peaking at week 15 followed by a decline at week 36. At this time point only one infant remained exclusively breastfed. The reduction in breastfeeding or the introduction of solid foods or formula could lead to a loss in MMc due to increased intestinal or systemic inflammation leading to rejection of maternal cells or due to the lack of continued transfer of maternal breastmilk cells. Distinguishing between these possibilities is challenging in humans but may be possible in future cohorts through improved characterization of the differences between breastmilk T cells and the maternal peripheral T cell repertoire. Of note, prior work has found that transplacentally acquired maternal cells traffic into fetal lymph nodes, whereas animal models suggest that breastmilk derived maternal cells traffic to the liver and lung (15,16), with possible differential impact on both priming and recall responses in the infant.

While the level of MMc we identified in most infants at any time point was similar to prior work (0-10/100,000 gEq) (5,56), in a proportion of infants we observed very high levels of MMc at select time points, such that up to 1% of cells appeared to be of maternal origin, many orders of magnitude higher than previously described. This observation raises many essential follow up questions about the origin, phenotype, and function of these cells. Specifically, we hypothesize that these discrete high levels of MMc may represent maternal cell proliferation in response to an exogenous stimulus (e.g. infection or vaccination in the infant), which should be evaluated in future studies.

In order to understand the functional consequence of MMc, we asked whether birth or concurrent MMc was associated with polyfunctionality of BCG CD4+ T cell responses in the infant. Polyfunctional T cells may be indicative of the quality of immune responses to vaccines and have been associated with better clinical prognosis during tuberculosis (**TB**), HIV, and other infections as compared to the absolute magnitude of T cell responses (37,57-61). We were particularly motivated to assess this outcome due to prior data suggesting that HIV exposed infants may mount less functional immune responses to BCG (20,62,63). We identified a positive correlation between birth, but not concurrent, MMc and the polyfuntionality of the T cell response to BCG during early infancy. Since our recruiting site was in Khayelitsha, Western Cape, South Africa, with one of the highest TB rates globally (64,65), we hypothesized that the polyfunctional cells we detected in infants might reflect maternal mycobacteria-responsive T cells. Notably, we did not find evidence that this was the case, suggesting that maternal cells present at birth may play an indirect role in shaping infants’ immune responses to BCG. This alternative possibility is consistent with prior work in a murine model of MMc, which showed that MMc modulates the development of the myeloid lineage, subsequently enhancing neonatal immunity to pathogenic challenge (32). These observations suggest that lower MMc in iHEU at birth may contribute to altered BCG vaccine responses in these infants.

Our study has a number of limitations. First, our samples were limited to maternal-infant pairs where there was a non-shared, non-inherited maternal-specific marker. However, there were no differences in the characteristics of the mother-infant pairs that could or could not be targeted for measurement of MMc. Secondly, at the time of recruitment, HIV viral loads and CD4+ counts were not routinely checked as part of prenatal care, so the relationship between viral load, CD4+ count, and MMc could not be evaluated; however, based on local and regional data (66,67) we anticipate that >90% of the women in this cohort were virologically suppressed at the time of delivery. Due to robust prenatal screening and high maternal ART coverage, there was only one infant infected with HIV in the InFANT cohort, and very few in the Western Cape in South Africa, precluding our ability to study this group. Furthermore, HIV infection and ART exposure is associated with higher rates of preterm birth (68–71), however, premature infants were not eligible to enroll in this study, and we may have had different findings had they been included. Finally, BCG response data was only available for a subset of the cohort, although there were no appreciable differences between those infants with and without data available.

To our knowledge, this is the first study to assess MMc in infants across the first year of life and relate their levels to infant T cell response to vaccination. Our findings highlight the as yet largely unexplored impact of this cross-generation graft of maternal cells on infant immunity. Furthermore, our findings provide insight into an additional potential mechanism that may contribute to altered immune responses in iHEU and the importance of maternal treatment to impact infant outcome. Future work should investigate the cellular phenotype and antigen-specificity of these inherited maternal cells.

## Materials and Methods

### Cohort

Data and samples were utilized from the Innate Factors Associated with Nursing Transmission (InFANT) study, a prospective birth cohort study conducted in Khayelitsha, Cape Town, South Africa funded by the Canadian Institute for Health Research (Co-PIs Jaspan, Gray). Mother-infant pairs were recruited at the Midwifery Obstetric Unit at Site B in Khayelitsha, Cape Town, South Africa from March 2014 to March 2018. Infants were followed from birth, at day 4-7, and at weeks 7, 15, and 36 of life. Voluntary counseling and HIV testing was done at the time of antenatal care registration. Both pregnant women living with HIV (WLHIV) and women not living with HIV were eligible for the study. Pregnant WLWH and their infants were provided with ART according to the current country-specific guidelines (72). Exclusive breastfeeding was advised to all mothers from delivery to at least 6 months. Infants born before 36 weeks of gestation and with birth weights lower than 2.4 kg were ineligible for the study. Further exclusion criteria included pregnancy or delivery complications as previously described (73). All infants who were classified as HIV exposed but uninfected were confirmed as HIV negative by PCR at delivery and at later time points according to prevention of vertical transmission guidelines (72). Routine vaccines were given to all infants according to the WHO’s Expanded Program on Immunization (EPI). Infants received intradermal Danish BCG strain (1331, Statens Serum Institute, Denmark) from April 2013 to January 2016 and thereafter Russian strain (BCG-I Moscow, Serum Institute of India, India). Both strains were given at 2 × 10^5 CFU/dose at birth. Maternal-infant pairs were included in the present study based on available paired maternal and infant samples (n=90) and informative typing for a maternal-specific allele with an assay available (n=58).

### Genomic DNA Extraction

Whole blood samples were collected at day one (birth) and weeks 7, 15, and 36 of life into Sodium-Heparin tubes. A 200 μl aliquot of the whole blood was lysed using FACS lysis buffer and subsequently fixed and stored at –80°C until genomic DNA (**gDNA**) extraction. The gDNA was extracted from fixed whole blood (**WBF**) samples from infants and mothers using QiaAMP DNA Mini Kits (Qiagen) with slight modifications to the manufaturer’s protocol. In brief, WBF samples were thawed and centrifuged at 16,000 rpm for 2 mins to pellet cells. The supernatant was discarded and the pellet re-suspended in 180 μL Qiagen Buffer ATL and 20 μL Proteinase K followed by incubation at at 56°C for one hour. This was followed by addition of 200 μL Buffer AL and 200 μL 100% ethanol and the entire sample lysate was transferred onto the QIAamp MinElute column. The gDNA was eluted using 200 μL molecular grade water which was allowed to incubate on the membrane for 10 mins at room temperature to increase DNA yield.

### Identification of Maternal-Specific Polymorphisms and Microchimerism Evaluation

Genomic DNA from maternal and infant granulocytes collected at week 36 was genotyped at HLA class II loci DRB1, DQA1, and DQB1 by direct sequencing (Scisco Genetics, Seattle, WA). Noninherited, nonshared HLA polymorphisms were identified that could be used to evaluate MMc (3,74). Maternal-infant pairs with non-informative HLA typing were typed at 4 non-HLA loci: *GSTT1, ATIII, TG*, and *TNN*, targeting insertion/deletion polymorphisms (31). The maternal-specific polymorphism identified for each maternal-infant pair was selectively amplified from WBF gDNA using a panel of previously developed qPCR assays (31,74,75). The limit of detection of each of the qPCR assays is one target genomic equivalent (**gEq**) in up to 60,000 background gEq per reaction well (76). DNA from each WBF was distributed across multiple reaction wells targeting a total gEq of 120,000 per sample. A calibration curve for the polymorphism-specific assay was included to quantify the amount of MMc in each well, and the microchimeric gEq was summed across all tested wells. Total gEq tested for each sample was determined by targeting the nonpolymorphic β-globin gene, and a β-globin calibration curve (human genomic DNA [Promega]) was concurrently evaluated on each plate. Level of MMc is presented as the total microchimeric gEq proportional to the total gEq tested for each sample.

### Maternal-infant HLA Class II compatibility score

To analyze the degree of HLA Class II relatedness for each maternal-infant pair, we generated a scoring system based on whether the infant’s paternal allele at each locus (DPA1, DPB1, DQA1, DQB1, DRB1) was a match to self from the mother’s perspective. A score of 0 at a single locus signified no match between the infant’s paternally inherited allele and either of the mother’s two alleles; a score of 1 signified a match between the infant’s paternally inherited allele and at least one of the mother’s alleles at that locus. For each pair, we summed the score at each locus to give an HLA compatibility score between 0-5 (0 signifies no HLA relatedness between infant’s paternally derived allele and mother’s alleles at any loci; 5 signifies a match at each locus between the infant’s paternally derived allele and at least one of the mother’s alleles).

### Whole Blood BCG stimulation Assay

A 12-hour whole blood assay was used to evaluate vaccine responses as previously described (77). Briefly, 250 μL of anticoagulated blood was incubated with BCG, media or PHA at 37°C within 1 h of blood draw. After 5 h, Brefeldin-A (Sigma Aldrich) was added and incubated for an additional 7 h. Thereafter, red blood cells were lysed followed by washing and staining with LIVE/DEAD^®^ fixable Violet stain (ViViD, ThermoFisher). The cells were cryo-preserved in liquid nitrogen (LN_2_).

### Cell Staining, Antibodies, and Flow Cytometry for BCG whole blood assay

Staining and flow cytometry were conducted as previously described (36). In brief, batched stored samples were thawed quickly and washed, incubated in BD PermWash and then stained with the antibody cocktail mix. After incubation, cells were washed and then resuspended in PBS for cell acquisition using a Beckton Dickinson LSRII flow cytometer (SORF model). The following monoclonal antibody-fluorochrome conjugates were used: IL-2-R-phycoerythrin (PE), CD8-V500, IFN-γ-Alexa Fluor-700, TNF-α-PE-Cy7, Ki67-Fluorescein isothiocyanate (FITC), all from BD, CD27 PE-Cy5, HLA-DR-Allophycocyanin-Cy7 (APC-Cy7), CD3-BV650 (BioLegend), CD4 PE-Cy5.5 (Invitrogen), CD45RA PE-Texas Red-X (Beckman Coulter). A minimum of 50,000 ViViD negative (viable) CD3 events were collected using FACS DIVA v6 software. Post-acquisition compensation and analysis was performed in FlowJo version 9 (FlowJo, LLC). **Supplementary Figure 2** shows the gating strategy employed. Measurable response to BCG was characterized by the polyfunctionality of the CD3+ CD4+ T cell response assessed by analyzing permutations of TNF-α, IL-2 and IFN-γ expression after stimulation using COMPASS (Combinatorial polyfunctionality analysis of antigen-specific T-cell subsets) (37). PFS and FS, which summarize the functional profile of each subject, were calculated from posterior probabilities as described (37).

### Stimulation, Cell Staining, and Cell sorting for detection of maternal mycobacterial-specific T cells

PBMC were isolated via density gradient centrifugation from infant whole blood. Available PBMC from infants who had detectible MMc at birth (in whole blood), and a COMPASS PFS>0.6 at the concurrent timepoint were selected for stimulation with BCG culture filtrate proteins. PBMC were thawed and treated with BD FastTimmune™ CD28/CD49d costimulatory antibodies (1:100, cat.# 347690), brefeldin A (1:1000, BioLegend), and BD GolgiStop™ (1:1500). PBMC were then stimulated with BCG culture filtrate proteins (100 μg/mL, BEI Resources), media, or PMA (0.05 μg/mL, Millipore Sigma) and ionomycin (1 μM, Millipore Sigma) and incubated at 37°C, 5% CO_2_ for 6 hours, upon which the reaction was stopped with the addition of EDTA (2 mM). Cells were treated with human Fc block (BD, 1:200) and then stained with LIVE/DEAD™ Fixable Aqua (ThermoFisher) and the following extracellular monoclonal antibody-fluorochrome conjugates: CD3-PE-Cy7 (BioLegend, cat.# 344816), CD4-BV421 (BioLegend, cat.# 300532), CD8-PerCP-Cy5.5 (BioLegend, cat.# 301032), CCR7-BV785 (BioLegend, cat.# 353230) and CD45RO-PE-CF594 (BD, cat.# 562299) or CD45RO-APC (BioLegend, cat.# 304210). Cells were washed and then resuspended in Cyto-Fast™ Fix/Perm Buffer (BioLegend), followed by washing in Cyto-Fast™ Perm Wash solution (BioLegend). Cells were then stained with intracellular monoclonal antibody-fluorochrome conjugates: IL-2-PE (BioLegend, cat.# 500307), TNF-α-FITC (BD, cat.# 554512), and IFN-γ-BV605 (BioLegend, cat.# 506542). Cells were washed, resuspended in cell staining buffer, and run on a BD FACSAria II™ Cell Sorter with single-stained AbC™ Total Antibody or ArC™ Amine Reactive Compensation Beads as compensation controls. To sort BCG-stimulated T cells into a polyfunctional population (positive for three or two of the cytokines IL-2, TNF-α, IFN-γ), a monofunctional population (positive for one cytokine), and a non-responsive population (negative for the three cytokines), cells were gated according to the scheme in **Supplementary Figure 3**. Genomic DNA was isolated from each sorted population, and MMc was quantified using qPCR assays targeting a maternal-specific marker, as described above.

### Statistics

Statistical analysis was performed using Stata 14 software (StataCorp LP) and R (version 4.0.3). Detection of MMc at birth was evaluated by logistic regression with adjustment for the total number of gEq tested for each subject. Level of MMc at birth was evaluated with negative binomial regression, accounting for the number of microchimeric gEq detected as well as the total number of gEq assessed in each sample (7,31,75,76,78,79).This approach accounts for the nonnormal distribution of MMc data as well as the large number of zeros (78). The output of this model is DRR, which can be interpreted as “X number of microchimeric gEq in group A for every one microchimeric gEq in group B.” Each covariate of interest was first tested individually in univariate models. For the adjusted model, we included HIV exposure and mode of feeding a priori as they were primary covariates of interest and included other covariates if they were associated with either detection or level of MMc at the P<0.1 level (infant sex, maternal-infant HLA Class II compatibility score, and gravidity).

Detection of MMc across infancy was assessed using generalized estimating equation (GEE) models clustered by individual, a binomial outcome structure, and an independent correlation matrix. Level of MMc across infancy was assessed using GEE models clustered by individual, a negative binomial outcome structure, and an independent correlation matrix. In addition to the covariates mentioned above, mode of feeding and infant age in weeks were considered for inclusion in the adjusted models.

The effect of MMc on the polyfunctionality of BCG vaccine responses (PFS) were assessed using GEE models clustered by individual, a gaussian outcome structure, and an independent correlation matrix. We have previously shown that iHEU infants have altered T cell functionality to BCG (20) and that the type of BCG strain used for vaccination contributes strongly to the magnitude and polyfunctionality of vaccine response elicited in infants (36). We therefore adjusted for BCG strain, HIV exposure, and infant age in our analyses.

### Study approval

The study was approved by the University of Cape Town Human Research Ethics Committee (Rec ref 285/2012) and the Institutional Review Board of Seattle Children’s Hospital (Protocol #15690). All mothers in the study were of consenting age and provided written informed consent for their and their infant’s participation.

## Author contributions

CB, BA, HBJ, SBK, CMG, JLN, and WEH conceived and designed the experiments. HBJ and CMG designed and recruited the InFANT cohort. CB, BA, AK, AHO, and AHA processed samples and performed wet lab experiments. CB, WH, XS and AK analyzed the data. CB, HBJ and WEH wrote the original draft of the manuscript. All authors reviewed and edited the manuscript.

## Acknowledgements

The study was funded by National Institutes of Health/University of Washington Center for AIDS Research grant AI027757 (WEH), Burroughs Wellcome Fund award number CAMS 1017213 (WEH), Global Health Research Initiative (GHRI) award number THA-118568 (HBJ, CGM), National Institutes of Health grants K08AI135072 (WEH), R21AI157821 (WEH), R01AI120714-01A1 (HJ), U01AI131302 (HBJ, CGM), and T32AI007509 (BA). We would like to thank the women and infants from Khayelitsha, Cape Town, South Africa for their participation in the InFANT cohort and recognize the clinic staff at Site B Khayelitsha and the InFANT clinical team for their care of study participants. We also acknowledge the Western Cape Department of Health and the Wellcome Centre for Infectious Diseases Research in Africa (CIDRI-Africa) for access to their facilities and equipment. We thank Katherine Guthrie, PhD for her assistance in selecting a statistical approach. Graphical abstract was generated using BioRender.com.

