## Supplementary Information for "Factors influencing maternal microchimerism throughout infancy and its impact on infant T cell immunity"

### Supplementary Tables

**Supplementary Table 1. Cohort characteristics stratified by informative maternal-specific marker**

| | **Informative & available assay (n=58)** | **Not informative or no available assay (n=32)** | **P value** |
|---|---|---|---|
| **Maternal age years, mean (sd)** | 27 (5.1) | 28 (4.5) | 0.313 |
| **Gravidity, n (%)** | | | 0.321 |
| Primigravidae | 12 (20.7) | 10 (31.3) | |
| Secondigravidae | 25 (43.1) | 9 (28.1) | |
| Multigravidae | 21 (36.2) | 13 (40.6) | |
| **Infant sex, female, n (%)** | 26 (44.8) | 19 (59.4) | 0.271 |
| **Birth weight, grams, mean (sd)** | 3,207 (416) | 3,114 (369) | 0.388 |
| **Gestational age at delivery, weeks, mean (sd)** | 39.0 (1.35) | 38.8 (1.06) | 0.480 |
| **Delivery method, vaginal, n (%)** | 58 (100) | 32 (100) | NA |
| **HIV exposed, n (%)** | 37 (63.8) | 18 (56.3) | 0.634 |
| **Maternal antiretroviral therapy** | | | 0.694 |
| Prior to pregnancy, n (%) | 20/37 (54.1) | 11 (61.1) | |
| Initiated during pregnancy, n (%) | 17/37 (45.9) | 7 (38.9) | |
| - Gestational age at initiation, weeks, mean (sd) | 21 (8.3) | 17 (7.7) | 0.389 |
| **BCG strain used for vaccination** | | | 0.274 |
| BCG-Denmark, n (%) | 39 (67.2) | 17 (53.1) | |
| BCG-Russia, n (%) | 19 (32.8) | 15 (46.9) | |

**Supplementary Table 2. Association between mode of feeding and detection and level of maternal microchimerism across infancy in a sensitivity analysis excluding week 36 timepoint.**

| | Detection of MMc across infancy (excluding week 36) | | | |
|---|---|---|---|---|
| | **OR (95% CI)** | **P value** | **Adj. OR (95% CI)** | **P value** |
| **Maternal HIV** | - | - | 0.68 (0.29-1.58) | 0.373 |
| **HLA Compatibility** | - | - | 1.46 (1.01- 2.11) | **0.041** |
| **Gravidity** | | | | |
| Primigravidae | - | - | REF | |
| Secondigravidae | | | 2.13 (0.68-6.60) | 0.675 |
| Multigravidae | | | 1.28 (0.40-4.09) | 0.192 |
| **Infant sex, female** | - | - | 1.42 (0.62-3.22) | 0.407 |
| **Exclusively breastfed** | 0.74 (0.34-1.62) | 0.446 | 0.90 (0.39-2.10) | 0.815 |
| **Infant age, weeks** | - | - | 1.04 (0.99-1.10) | 0.114 |
| | **Level of MMc across infancy (excluding week 36)** | | | |
| | **DRR (95% CI)** | **P value** | **Adj. DRR (95% CI)** | **P value** |
| **Maternal HIV** | - | **-** | 0.32 (0.12-0.81) | **0.017** |
| **HLA Compatibility** | - | **-** | 2.93 (2.03-4.23) | **<0.001** |
| **Gravidity** | | | | |
| Primigravidae | - | **-** | REF | |
| Secondigravidae | | | 0.92 (0.28-3.09) | 0.897 |
| Multigravidae | | | 2.07 (0.41-10.6) | 0.380 |
| **Infant sex, female** | - | **-** | 4.31 (1.32-14.1) | **0.015** |
| **Exclusively breastfed** | 11.5 (3.13-42.3) | **<0.001** | 2.73 (0.92-8.84) | 0.068 |
| **Infant age, weeks** | - | **-** | 1.18 (1.06-1.32) | **0.003** |

Detection of maternal microchimerism (MMc) across infancy was analyzed using generalized estimating equation (GEE) models with binomial distribution, controlling for total genomic equivalents (gEq) assessed. Level of MMc across infancy was analyzed using GEE models with negative binomial distribution accounting for both microchimeric and total gEq. Adj.: adjusted, DRR: detection rate ratio, OR: odds ratio.

### Supplementary Figures

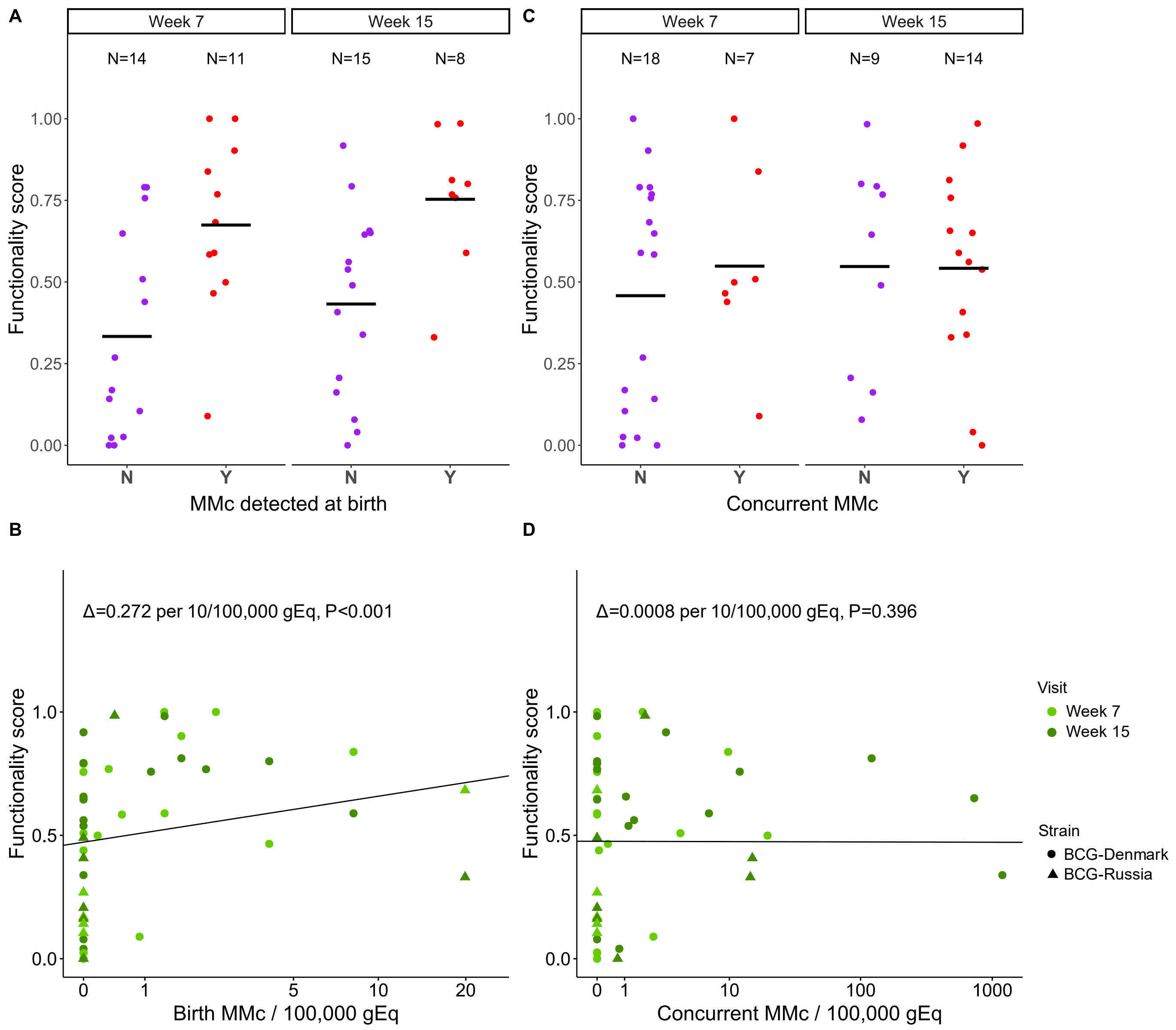


**Supplementary Figure 1. Maternal microchimerism at birth is positively associated with T cell responses to BCG vaccination.** (A) Functionality score (FS) at week 7 and week 15 of life by detection of maternal microchimerism (MMc) at birth (N=No (purple), Y=Yes (red)). (B) Association between FS at week 7 (light green) and week 15 (green) of life with level of MMc per 100,000 genomic equivalents (gEq) at birth. Delta represents the adjusted effect size per 10/100,000 gEq. (C) FS at week 7 and week 15 of life by detection of MMc at the concurrent time point (Y=Yes (red), N=No (purple)). (D) Association between FS at week 7 (light green) and week

15 (green) of life by level of MMc per 100,000 gEq at the concurrent time point. Delta represents the adjusted effect size per 10/100,000 gEq.

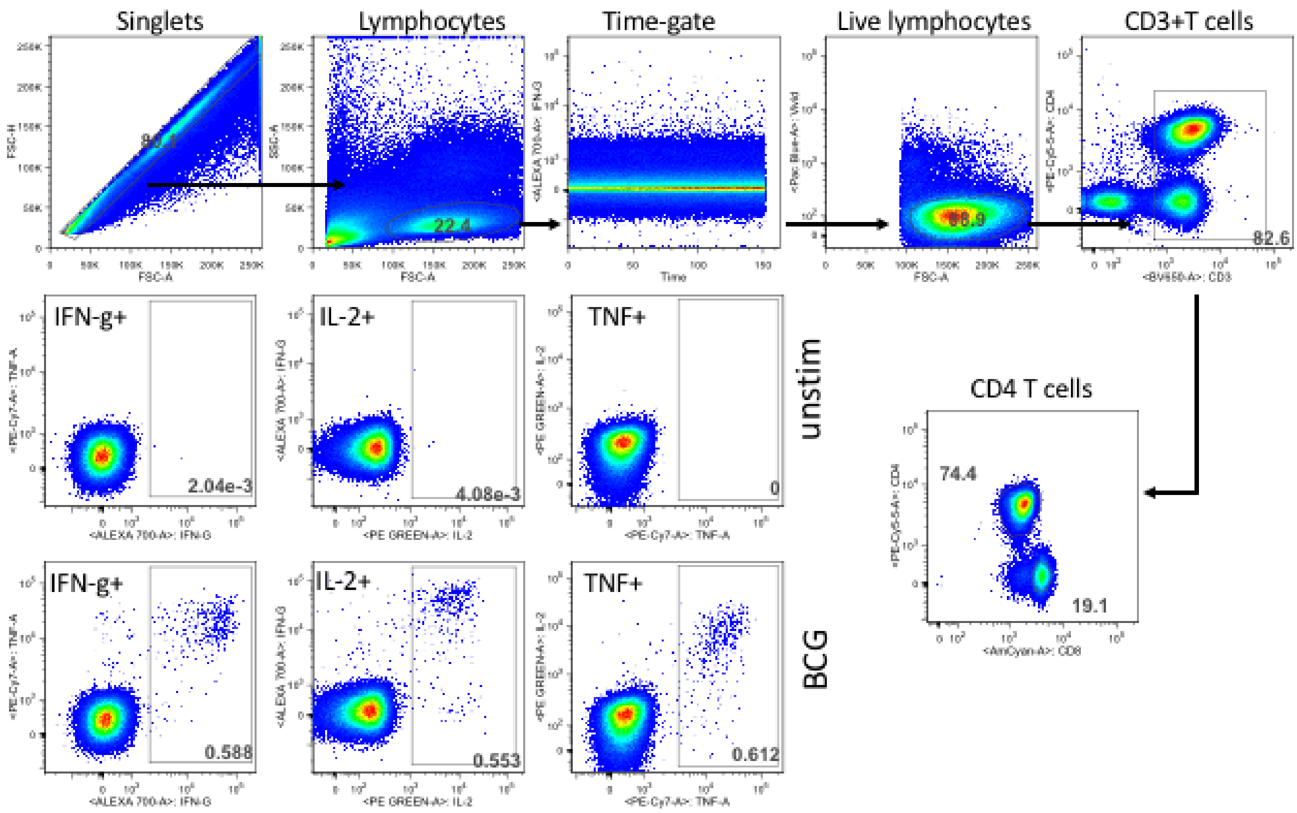


**Supplementary Figure 2. Flow cytometry gating strategy for quantifying antigen specific responses.** Total cytokine responses were defined based on any combinations of live CD4+ T cells expressing any of IL-2, TNF-α or IFN-γ total cytokine population from the parent CD4+ population from antigen-stimulated samples.

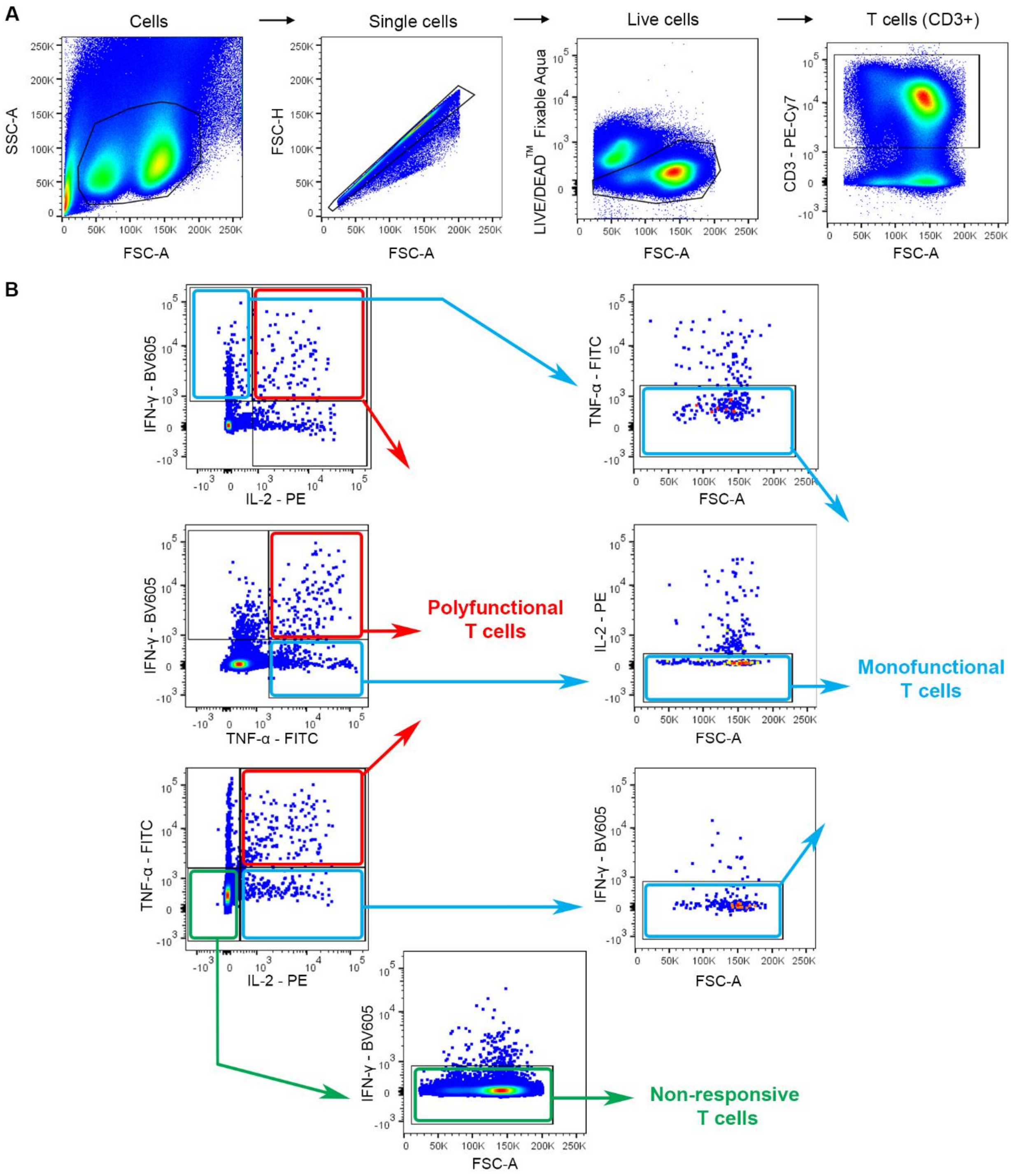


**Supplementary Figure 3. Flow cytometry gating strategy for sorting infant PBMC stimulated with BCG culture filtrate proteins.** (A) Gating strategy for T cells (CD3+). (B) T cells (CD3+) were sorted into polyfunctional, monofunctional, or non-responsive groups. Polyfunctional T cells were defined as T cells that were triple-positive for IFN-γ, TNF-α, and IL-2 or double-positive for any two of these cytokines. Monofunctional T cells were defined as T cells that were single positive for IFN-γ, TNF-α, or IL-2. Non-responsive T cells were defined as T cells that were triple negative for IFN-γ, TNF-α, and IL-2.
